## Supplementary figures and images for "TCR transgenic clone selection guided by immune receptor analysis and single cell RNA expression of polyclonal responders"

### Supplemental Figure 1

#### Supplemental figure 1

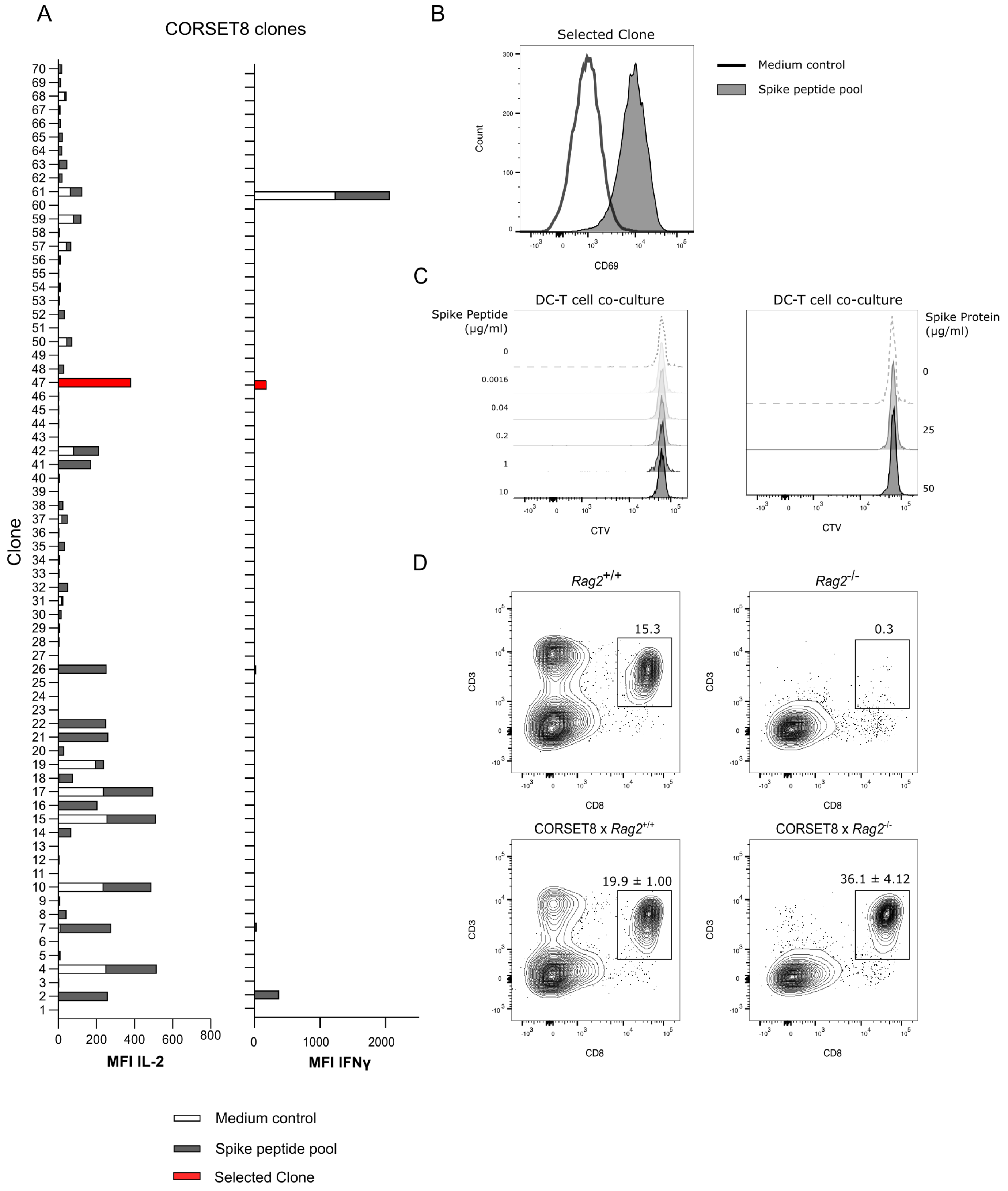

### Supplemental Figure 2

#### Supplemental Figure 2

Gating strategy: Non debris -> Single cells -> Viable cells

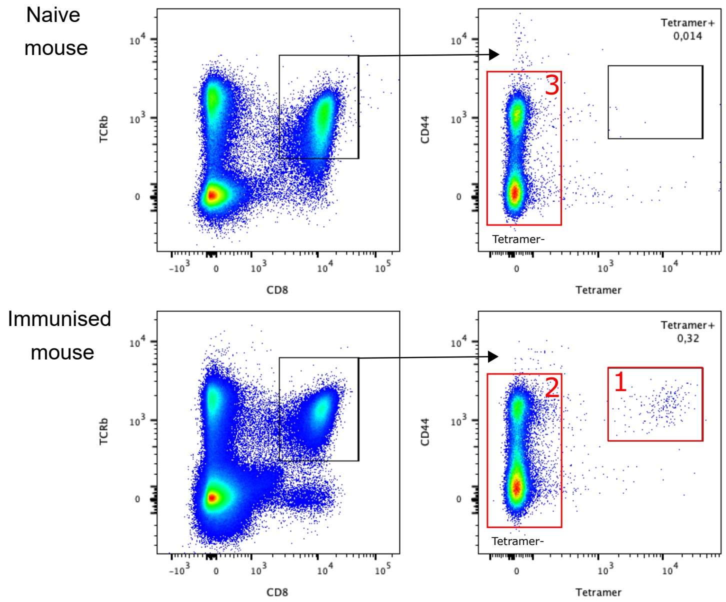
